# PEGPH20, a PEGylated Human Hyaluronidase, Induces Radiosensitization by Reoxygenation In Pancreatic Cancer Xenografts. A Molecular Imaging Study

**DOI:** 10.1101/2022.01.11.475894

**Authors:** Tomohiro Seki, Yu Saida, Shun Kishimoto, Jisook Lee, Yasunori Otowa, Kazutoshi Yamamoto, Gadisetti VR Chandramouli, Nallathamby Devasahayam, James B. Mitchell, Jeffery R. Brender, Murali C. Krishna

## Abstract

PEGylated human hyaluronidase (PEGPH20) enzymatically depletes hyaluronan, an important component of the extracellular matrix, in tumors. The resultant improvement in vascular patency and perfusion has been shown to increase the delivery of therapeutic molecules. We show that PEGPH20 also improves the efficacy of radiation therapy in a human pancreatic adenocarcinoma BxPC3 mouse model overexpressing hyaluronan synthase 3 (BxPC3-HAS3) while exerting little effect on the corresponding wild type tumors. Mice overexpressing HAS3 developed fast growing, radiation resistant tumors that became rapidly more hypoxic as time progressed. Treatment with PEGPH20 increased survival times when used in combination with radiation therapy, significantly more than either radiation therapy or PEGPH20 alone. Radiosensitization in BxPC3-HAS3 tumors was attributed to an increase in local pO_2_ as studied by by EPR imaging. No effect on survival, radiation treatment, or pO_2_ was seen in wild type tumors after PEGPH20 treatment. Dynamic contrast enhanced (DCE) MRI and MRI based blood volume imaging showed improved perfusion/permeability and local blood volume, respectively, in BxPC3-HAS3 tumors after PEGPH20 treatment, accounting for the increase in tumor oxygenation. Photoacoustic imaging indicated immediate changes in tumor oxygenation after treatment. Metabolic MRI using hyperpolarized [1-^13^C] pyruvate suggested a metabolic shift towards decreased glycolytic flux after PEGPH20 treatment. In summary, the results showed that PEGPH20 may be useful for radiosensitization of pancreatic cancer but only in the subset of tumors with substantial hyaluronan accumulation and the response of the treatment may potentially be monitored non-invasive imaging of the hemodynamic and metabolic changes in the tumor microenvironment.

## Introduction

Pancreatic ductal adenocarcinoma (PDAC) is characterized by an intense desmoplastic response which can significantly influence the tumor microenvironment. The dysregulation of growth signaling during the development of PDAC leads to a high expression of growth factors in the surrounding stroma(1,2) leading to an increase in fibroblast growth and creating a dense, tangled extracellular matrix (ECM) that both suppresses the convective currents responsible for the transport of large macromolecules and acts as physical barrier to the diffusion of smaller ones.(3,4) The ECM is largely composed of meshwork of collagens and elastin embedded in a matrix of hyaluronan(HA)(5), a long polysaccharide formed of repeating disaccharide units of *N*-acetyl-D-glucosamine and D-glucuronic acid.(6) While hyaluronan is a minor component of the ECM matrix by weight, it has received substantial research focus due to the outsized role it plays in hindering transport through the ECM.(7) Hyaluronan accumulates water at its interface,(8) forming a hydrogel of bound water molecules(9) that forms viscous barrier to diffusion within the ECM.(4) Electrostatic repulsion from the negative charges on hyaluronan also tends to expand the ECM network,(9) possibly creating a solid compressive stress on the blood vessels within.(10,11) The accumulation of water by hyaluronan also creates a hydraulic pressure (the interstitial fluid pressure) that resists permeation of therapeutics out of capillaries.(7,9,12)

Hyaluronan is highly expressed in most PDAC tumors.(13,14) PDAC is largely resistant to chemotherapy,(15,16) a factor that is not captured by many mouse xenograft models.(17) High HA expression is correlated with poor prognosis in patients with PDAC,(18) which may be associated with the poor penetration of anti-cancer drugs, compromising the therapeutic effect of chemotherapy.(19,20) Thus, depleting the ECM including HA may be a potentially viable method to improve the treatment efficacy of cancer therapies of solid tumors.

Towards this end, a pegylated version of human hyaluronidase (PEGPH20) that enzymatically depletes intra-tumor HA and re-expands the intratumor vasculature was advanced as a chemotherapy adjuvant. Preclinical trials of PEGPH20 showed increased delivery of small molecule chemotherapeutic molecules into PDAC and other tumors.(13,21-23). Based on these results, PEGPH20 was advanced into phase 2 clinical trials in combination with gemcitabine and nab-paclitaxel for pancreatic ductal adenocarcinoma where it demonstrated a favorable safety profile and promises of enhanced efficacy.(24) Unfortunately, results from the Phase III trials were less impressive. PEGPH20 in combination with nab-paclitaxel/gemcitabine for metastatic pancreatic cancer reduced tumor burden but had no impact on progression free or overall survival.(25) A similar Phase II trial where patients were not preselected for HA expression was stopped early for lack of efficacy and adverse events.(26)

The reasons for the failure of the HALO301(27) and SWOG1313(26) trials are as yet unexplained but may stem from a general resistance to chemotherapy from the suppression of nucleoside transporters necessary for gemcitabine transport during the epithelial-to-mesenchymal transition in high grade PDAC tumors.(28,29) The effect of PEGPH20 on other treatment modalities such as radiation an immunotherapy is largely unexplored. The elevated pressure from HA expansion can cause vascular collapse and decreased blood perfusion in the tumor microenvironment, which can lead in turn to local hypoxia (13,23). We show here using multimodal imaging techniques to comprehensively examine changes in tumor physiology and metabolism in response to the treatment with PEGPH20 that in mouse models overexpressing hyaluronan synthase 3, PEGPH20 treatment leads to a rapid increase in blood vessel patency and transport that raises pO_2_ levels within the tumor. The increase in oxygen concentrations leads to a reduction in hypoxia and shifts metabolism away from a primarily aerobic glycolysis to the TCA cycle. In this environment, PEGPH20 treatment led to a significant increase in the ability of radiation therapy to suppress tumor growth and improvement in survival. The imaging techniques are non-invasive and can be serially applied to examine physiologic and metabolic profiles in response to therapies which impact the tumor microenvironment. The data therefore not only provides a direct rationale for PEGPH20 use in radiotherapy, but may also help optimizing the regimen and timing of its use.

## Materials and Methods

### Mice and tumors

PEGPH20 and BxPC3 PDAC (human pancreatic adenocarcinoma) cells lines transduced with hyaluronan synthase 3 (HAS3) were provided by Halozyme Inc.. The BxPC3 wild type cell line was purchased from ATCC. The identity of both cell lines was verified by CellCheck9 from IDEXX BioAnalytics. Athymic nude mice were inoculated with 2 x 10^6^ BxPC3-HAS3 or BxPC3-wild tumor cells adjacent to the right tibial periosteum. For PEGPH20 treatment, mice bearing an approximately 600 mm^3^ tumor were i.v. injected with 1 mg/kg of PEGPH20 on day 0 and day 3. Tumor bearing mice in the control group were injected with same amount of API buffer. All imaging experiments except for photoacoustic imaging were performed at the the same times pre-treatment (before the first injection on day 0) and post-treatment (3 h after the 2nd PEGPH20 injection on day 3). Photoacoustic imaging was performed for 1 minutebefore and 20 minutes after the bolus injection of PEGPH20 (1 mg/kg or 10 mg/kg) to monitor temporal changes in local hemoglobin saturation. For radiotherapy, approximately 600 mm^3^ BxPC3-HAS3 or BxPC3-wild tumor bearing mice were injected i.v. with 1 mg/kg of PEGPH20 3h prior to radiotherapy. Radiotherapy was performed at 2.5 Gy. PEGPH20 and radiotherapy were performed 5 times in total, given once every three days (q3d).

### EPRI

Technical details of the EPR scanner and oxygen image reconstruction were described previously (30). Parallel coil resonators tuned to 300MHz were used for EPRI. After the animal was placed in the resonator, an oxygen-sensitive paramagnetic trityl radical probe, OX063 (1.125 mmol/kg bolus), was injected intravenously under isoflurane anesthesia. The free induction decay (FID) signals were collected following the radiofrequency excitation pulses (60 ns) with a nested looping of the x, y, and z gradients, and each time point in the FID underwent phase modulation, enabling 3D spatial encoding. Because the T2* of OX063 is strongly affected by the presence of oxygen, it is possible to generate a series of T2* maps, i.e., EPR line width maps, which linearly correlate with the local concentration of oxygen if the concentration of Ox063 is low enough to avoid the contribution of self-line broadening, and allow pixel-wise estimation of tissue pO_2_. The repetition time was 8.0 μs. The number of averages was 4,000. After EPRI measurement, corresponding anatomic T2-weighted MR images were collected with a 1T scanner (Bruker BioSpin MRI GmbH).

### DCE-MRI of Gd-DTPA

DCE-MRI studies were performed on a 1 T scanner (Bruker BioSpin MRI GmbH). T1-weighted fast low-angle shot (FLASH) images were obtained with TR = 117.2 ms; TE = 6 ms; flip angle = 30°; two slices; FOV = 28 x 28 mm; matrix = 64 x 64; 15-second acquisition time per image; 45 repetitions; number of average = 2. Gd-DTPA solution (4 mL/g of body weight of 50 mmol/L Gd-DTPA) was injected through a tail vein 1 minute after the start of the dynamic FLASH sequence. To determine the local concentrations of Gd-DTPA, T1 maps were calculated from four sets of Rapid Imaging with Refocused Echoes (RARE) images obtained with TR = 300, 600, 1000, and 2,000 ms, with the acquisitions being made before running the FLASH sequence.

### Photoacoustic imaging

Tumors were imaged with the VisualSonics Vevo®LAZR System (FUJIFILM VisualSonics Inc.) using a 21-MHz linear array transducer system (central frequency) integrated with a tunable nanosecond pulsed laser. The tumor area in the sagittal plane of the leg was determined manually from concurrently acquired ultrasound images. For O_2_ status assessments, photoacoustic images in the tumor area were collected with OxyHemo-Mode (wavelength 750 nm/850 nm) every 3 s for 25 min. Oxygen saturation of hemoglobin (sO_2_) in the tumor area were calculated using OxyZated™ tool.

### Blood volume imaging

MRI scanning was conducted on a 3-T scanner controlled with ParaVision 6.0 Bruker BioSpin MRI). For blood volume (BV) calculation, spoiled gradient echo sequence images were collected before and 5 minutes after injection of ultra-small superparamagnetic iron oxide (USPIO) contrast (1.2 μL/g of body weight). The imaging parameters included the following: FOV = 28 x 28 mm; matrix = 192 x 192; echo time (TE) = 3.5 ms; TR = 200 ms; and number of averages = 12. The percentage tumor BV was estimated as described previously (31).

### Hyperpolarized ^13^C MRI

Details of the hyperpolarization procedure were reported previously (31,32). Briefly, [1-^13^C] pyruvic acid (30 μL) containing 15 mmol/L Ox063 and 2.5 mmol/L of the gadolinium chelate ProHance (BraccoDiagnostics) was polarized in Hypersense DNP polarizer (Oxford Instruments). After the polarization reached 80 %of the plateau value, the hyperpolarized sample was rapidly dissolved in 4.5 mL of a superheated alkaline buffer consisted of 40 mmol/L HEPES, NaOH, and 100 mg/L EDTA. Hyperpolarized [1-^13^C] pyruvate solution was rapidly injected intravenously through a catheter placed in the tail vein of each mouse (12 mL/g body weight). Hyperpolarized ^13^C MRI studies were performed on a 3 T dedicated MR Solutions animal scanner (MR SOLUTIONS Ltd., Boston, MA) using a 17 mm home-built ^13^C solenoid coil placed inside of a saddle coil tuned to ^1^H frequency. Both ^1^H and ^13^C were tuned and matched and anatomical image was taken after shimming on proton. ^13^C spectra was acquired every 1 s for 240 s from the whole leg with a tumor. The repetition time, spectral width, flip angle, and number of averages were 1000 ms, 3300 Hz, 10°, and 1, respectively.

### Histological assessment

Tumor tissues were excised 1 hour after intravenous injection of pimonidazole (60 mg/kg, Hypoxyprobe Inc.), frozen with Tissue-Tek O.C.T. compound (Sakura Finetek USA Inc.) by ultracold ethanol and sectioned (10 mm). After fixing with 4 % paraformaldehyde, sections were treated with cold acetone for 15 minutes. After blocking nonspecific-binding sites on sections with Protein Block Serum-Free reagent (Dako North America Inc.) for 30 minutes, the slides were covered by pimonidazole antibody (Hypoxyprobe-1 Omni Kit, Hypoxyprobe Inc.; 1:250) overnight at 4 C. The sections were then incubated with Alexa Fluor 488 secondary antibody for 1 h at room temperature, before being mounted with Prolong Gold antifade reagent with DAPI (Invitrogen). For γH2AX staining, anti-γH2AX antibody (Abcam, Inc.; 1:250) was used, followed by same staining procedures as above. The stained slides were scanned using a BZ-9000 microscope (Keyence), and the immunostain-positive area was quantified using ImageJ software (downloaded from https://imagej.nih.gov/ij/).

### Statistical analysis

The significance of the differences between pre-treatment and post-treatment within the same treatment group was analyzed using the paired *t*-test. The significance of the differences between groups was analyzed using the Student’s *t*-test. Kaplan-Meier curves were constructed for the survival of mice; the differences between multiple groups were identified using the Bonferroni test. A two-tailed *p* value of 0.05 was considered significant.

## Results

### PEGPH20 and radiation therapy work together to inhibit the tumor growth of BxPC3-HAS3 tumor

In a preclinical setting, it has been suggested that the tumor ECM as a whole can induce resistance and recurrence to radiation therapy in different cancers,(33,34) but the specific effect of HA on radiosensitization remains untested. To determine the radio-sensitizing effect of depleting HA by PEGPH20, we compared the effect of radiation therapy and PEGPH20 treatment on BxPC3-HAS3 mice xenografts which overexpress HA and wild type BxPC3 tumors (BxPC3-WT) which do not (Figure 1).

**Figure 1.**
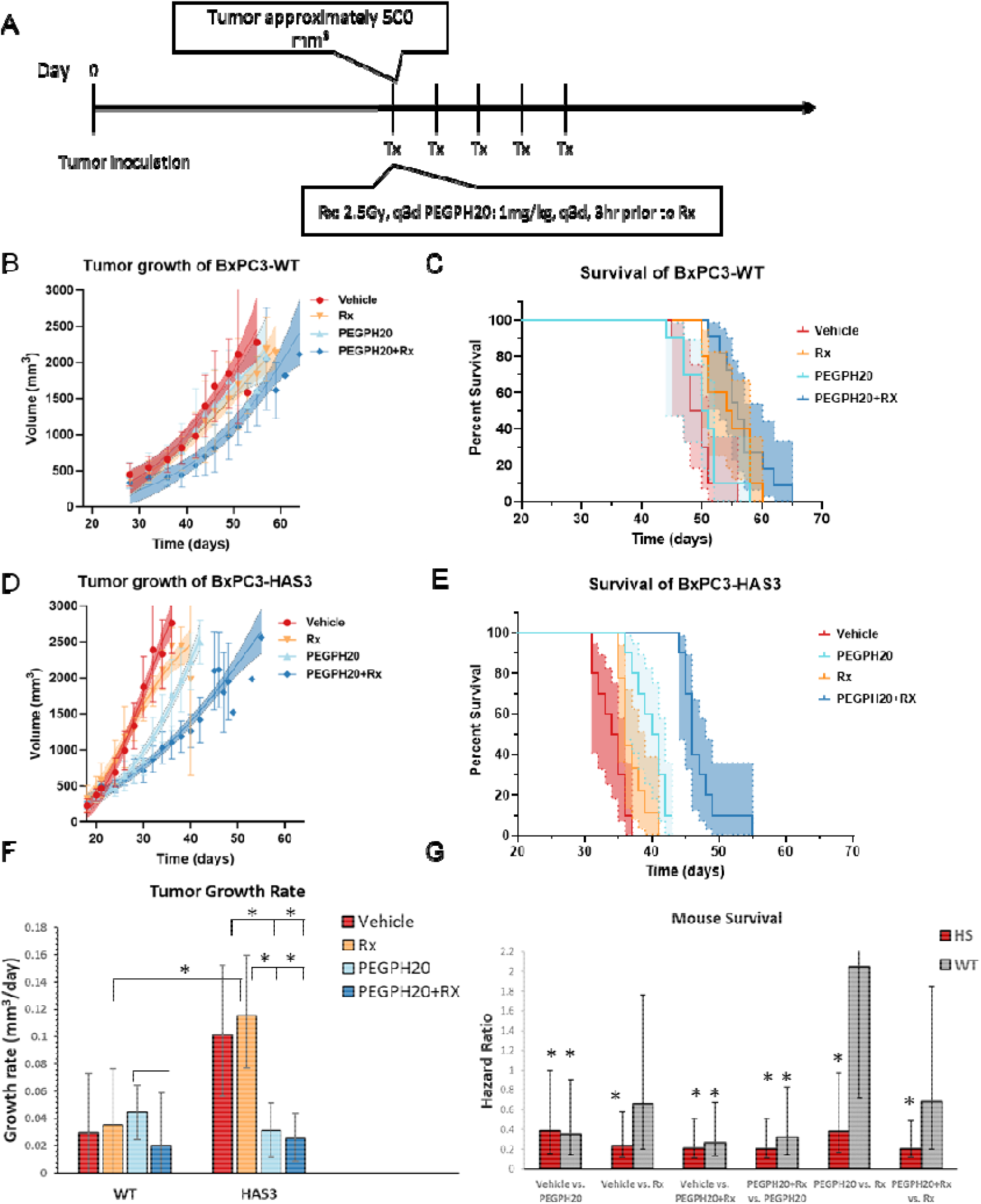
Tumor growth and survival of BxPC3-HAS3 and BxPC3-wild tumors in response to radiation and PEGPH20 therapy. **A** Treatment plan. Treatment was initiated when the tumor size reached approximately 500 mm3 and consisted of a single dose of 2.5 Gy radiation and/or 1mg/kg PEGPH20 given every 3 days. **B** Growth kinetics of BxPC3-wild type tumors **C** Kaplan-Meier survival curve for BxPC3-wild type tumors (n = 10 per group). Survival refers to the time before reaching the maximally allowed tumor volume of 2,500 mm^3^. Asymmetric 95% CIs (shaded region) were determined by the profile method. **D** Growth kinetics of BxPC3-HAS3 tumors **E** Kaplan-Meier survival curve for BxPC3-wild type tumors (n = 10 per group). **F** Growth rates estimated from a Gompertz logistic growth model of the data in B and D. **G** Hazard ratios of the treatment effect calculated (lower numbers favor the second treatment listed)

Compared to wild type tumors, BxPC3-HAS3 tumors grew faster in the absence of therapy (Figure 1B, C and F, red) and were associated with lower survival times (Figure 1D,E and G), in line with previous preclinical(35) and clinical(18) results. Both BxPC3-HAS3 and BxPC3-WT tumors were relatively resistant to radiation monotherapy when treatment was initiated when the tumor size reached approximately 500 mm^3^ (Figure 1, orange). Resistance to radiation monotherapy was greater in BxPC3-HAS3, which showed almost no effect of radiation monotherapy on survival times (median survival time 36 days vs. 35 days in the control group, HR=0.7, 95% CI 0.1 - 1.0). Wild type BxPC3 were significantly more sensitive to radiation (median survival time 55 days vs. 49 days in the control group, HR=0.23 vs. control, 95% CI=0.11-0.35). Depletion of HA by PEGPH20 had little effect on wild type tumors; PEGPH20 failed to slow tumor growth and did not lead to an increase in survival time (median survival time 50.5 days vs. 49 days in the control group, HR=0.66, 95% CI= 0.27 – 1.61). PEGPH20 was considerably more effective than in BxPC3-HAS3 tumors. Tumor growth was 69% slower than in the control (95% CI=62 – 88%) and survival was extended from a median of 34.5 days to 40.5 days (HR=0.21, 95% CI 0.10 - 0.30). As a monotherapy, PEGPH20 was more effective than radiation in BxPC3-HAS3 (HR=0.38, 95% CI 0.21 – 0.59) but less effective in WT tumors (HR=2.1, 95% CI=1.3 – 3.4).

Radiation therapy worked in concert with PEGPH20 in both WT and BxPC3-HAS3 tumors to suppress tumor growth and improve survival times. In BxPC3-HAS3 tumors, the combination treatment significantly suppressed tumor growth and improved survival compared to either treatment alone (HR= 0.20, 95% CI=0.08-0.29 vs. radiation alone and HR=0.21, 95% CI=0.09-0.3 vs PEGPH20 alone). In WT tumors, the effect was less and did not reach statistical significance when compared to radiation alone (0.68, 95% CI=0.48 vs. 1.17). Overall, the results suggested that the involvement of excessive accumulation of HA is associated with an aggressive, radiation-resistant phenotype that is vulnerable to PEGPH20 treatment.

### Changes in intratumor pO2 induced by PEGPH20

The dense and rigid extracellular matrix formed by HA accumulation can encapsulate clusters of tumor cells, resulting in a barrier that constricts blood vessels and reduces the available oxygen supply.(36) Since oxygen sensitizes the tumor to radiation therapy, we hypothesized that the the radiosensitization of PEGPH20 treatment was conferred by improved tumor oxygenation within the tumor after PEGPH20 treatment. To test this hypothesis, we measured pO_2_ levels in tumor-bearing mice treated with either the vehicle or PEGPH20 by EPR oximetry imaging, which in contrast to other methods, directly reports on the partial oxygen pressure in tissues that can be perfused with the spin probe OX063 to a detectable degree.

Most tumors were highly hypoxic, as shown by the representative pO_2_ maps of a BxPC3-HAS3 xenograft before and 3 hours after treatment in Fig. 2A. In the absence of treatment, median pO2 levels within the tumor (pink line, Fig. 2A) decreased in the 3 day period between scans, especially in the interior of the tumor (Fig. 2A, histogram 2B). This change was much more pronounced in the BxPC3-HAS3 xenografts (−12.2 ± 3.5 %, *p* = 0.0155, Figs. 2C and D) compared to the BxPC3-wild type (Figs. 2E and F) where the change over the 3 day period was minor and not statistically significant.

**Figure 2.**
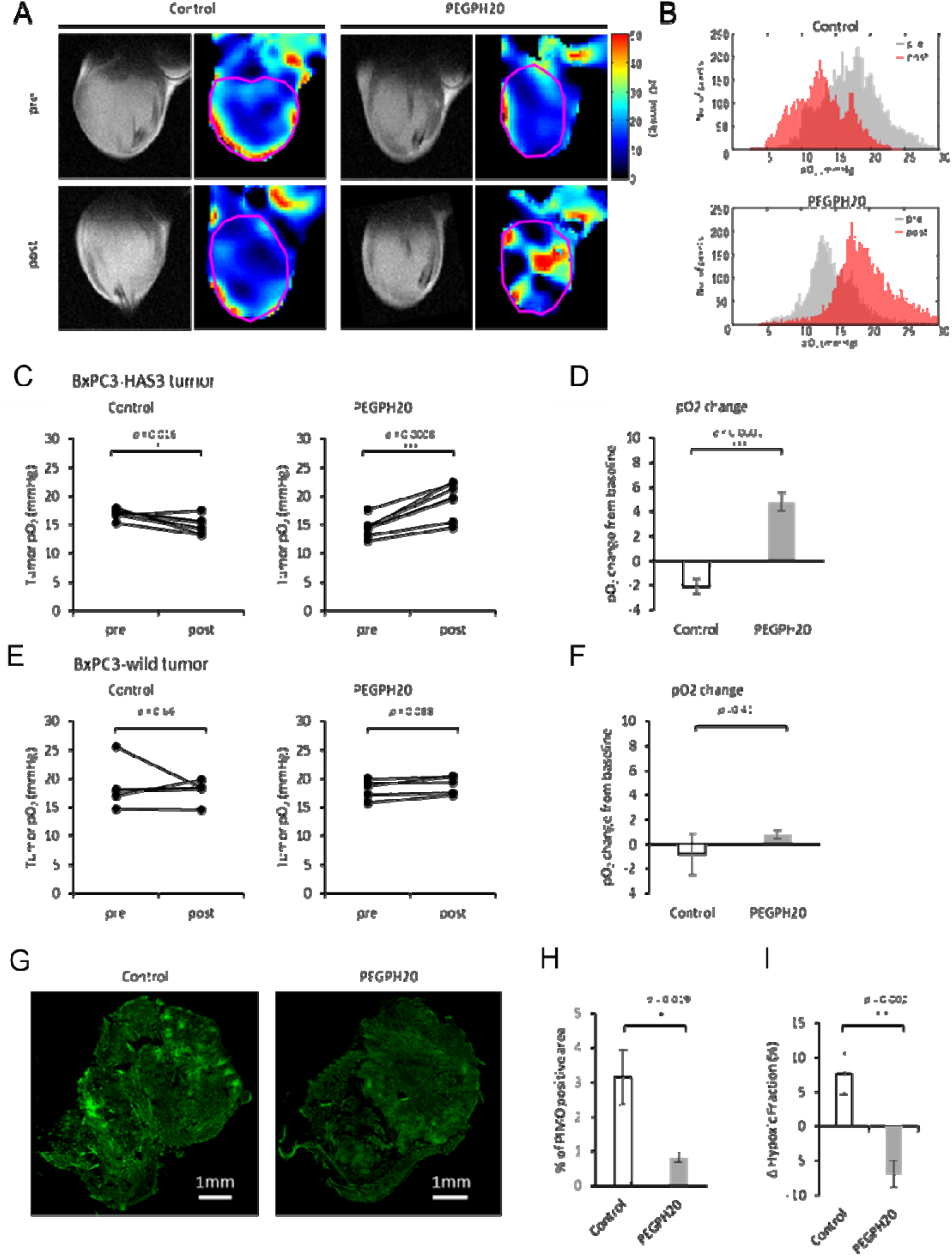
EPR imaging showed increased pO_2_ after PEGPH20 treatment. EPR imaging was performed in BxPC3-HAS3 tumor bearing mice treated with control buffer or PEGPH20 (n = 7 each group) before (before the first injection on day 0) and after treatment (3 h after the 2nd Ab injection on day3). **A**, Representative oxygen map obtained by EPR imaging and T2-weighted anatomical image. **B**, Histograms of pO_2_ distribution within the tumor of each group. **C and E,** The mean pO_2_ of each tumor before and after treatment are shown by treatment group. Individual values are shown. **D and F,** pO_2_ change (%) from the baseline (pretreatment) of each group. Data are shown as mean ± SE. **G and H,** Histological assessment of pimonidazole staining. BxPC3-HAS3 tumors treated with control buffer or PEGPH20 were surgically resected 3h after the 2^nd^ Ab injection on day3. Frozen sections of tumors were stained for exogenous hypoxia marker pimonidazole (pimo). **G,** Representative pimo staining of each group (scale bar = 1mm). **H,** Quantification of pimo positive area. **I,** Comparison of ΔHF10 between control and PEGPH20 groups evaluated by EPR imaging. Data are shown as mean ± SE (n = 5 per group). Statistical significance between groups was determined by paired *t*-test for **C** and by Student’s *t*-test for D and F. \**p* < 0.05, \*\**p* < 0.01, \*\*\**p* < 0.001 \*\*\*\**p* < 0.0001.

PEGPH20 treatment led to a substantial reoxygenation effect in BxPC3-HAS3 xenografts (*p* < 0.0001, Figs. 2C and D) after two i.v. injections of 1 mg/kg three days apart. While tumor pO_2_ levels decreased in the 3 day period between injections in the absence of treatment (−12.2 ± 3.5 %, *p* = 0.0155), they increased by a median 32.5 ± 5.0 % (*p* = 0.0008) after PEGPH20 treatment. Reoxygenation of BxPC3-HAS3 xenografts after PEGPH20 treatment was also supported by immunohistochemical evaluation by pimonidazole, which binds to thiol-containing proteins specifically in hypoxic cells where pO_2_<10 mmHg. Figure 2G shows representative images of whole tumor treated with or without PEGPH20 stained with pimonidazole. The hypoxic fraction stained by pimonidazole in vehicle-treated tumors was more than 3-fold that of PEGPH20 treated tumors (Fig. 2H). A similar decrease in hypoxic fraction with pO_2_ levels <10 mmHg (ΔHF10) between control and PEGPH20 treated tumors was also observed after PEGPH20 treatment in EPR oximetry maps (7.6% increase in the control group compared to a 6.9% decrease in PEGPH20 group) (Fig. 2I).

PEGPH20 did not lead to a reoxygenation effect in the absence of HA overexpression. In BxPC3-wild type tumors, PEGPH20 group only slightly improved pO_2_ levels and the difference between the treatment and control groups was not statistically significant (Fig. 2E and 2F). These findings suggest that HA overexpression leads progressively to more severe hypoxia and HA depletion by PEGPH20 treatment improves oxygenation and significantly decreases the severely hypoxic region where radiation therapy is ineffective.

### Changes in Intratumor perfusion induced by PEGPH20

Reoxygenation is a complex effect that can have many causes, and can result from oxygen demand decreasing due necrosis/apoptosis within the tumor or an increase in blood supply. We hypothesized a ECM dense in HA exerts pressure on neighboring structures resulting in elevated IFP, leading to a collapse of blood vessels and impaired blood transport. Decreasing HA within the ECM should decrease the tumor IFP and restore perfusion pressure and open micro-vessels and capillaries. To test this theory, tumor perfusion changes in response to PEGPH20 treatments were non-invasively examined by DCE-MRI using the Toft model. In this model, permeability changes are characterized by the influx forward volume transfer constant (K^trans^) from plasma into the extravascular-extracellular space (EES), which reflects the sum of all processes (predominantly blood flow and capillary leakage) that determine the rate of gadolinium influx from plasma into the EES. We examined BxPC3-HAS3 tumors treated with either vehicle or PEGPH20 (1 mg/kg) before and after the treatment (3 h after the 2nd PEGPH20 injection on day3). Figure 3A and Figure 3B show representative images and kinetics of intratumor Gd-DTPA concentration in tumors treated with buffer and PEGPH20. In the tumor treated with vehicle, only a minor change was observed in Gd-DTPA accumulation and its kinetics (Fig. 3A, 3B, upper panels). Some mice showed a decrease in intratumor perfusion, which was assumingly caused by the tumor progression during the 3 days of experiments. In contrast, Gd-DTPA enhancement increased and infiltrated even the center of the tumor after PEGPH20 treatment where enhancement was limited to the peripheral region before treatment (Fig. 3A, lower panels). The rapid increase in Gd-DTPA concentration in the first few minutes indicates improved perfusion after PEGPH20 treatment (Fig. 3B, lower panel). Figure 3C shows K^trans^ changes after the treatment. The mean K^trans^ decreased from 0.0126 ± 0.0021 to 0.0068 ± 0.0013 in the control group, while it increased from 0.0135 ± 0.0017 to 0.0238 ± 0.0048 in PEGPH20 group (*p* = 0.0358, Fig. 3C). The % change in K^trans^ was significantly higher in PEGPH20 group relative to the control (*p* = 0.0214, Fig.3D). Interestingly, perfusion was not improved in all tumors in the treated group, indicating some heterogeneity in reoxygenation possibly due to different levels of hyaluronan expression.

**Figure 3.**
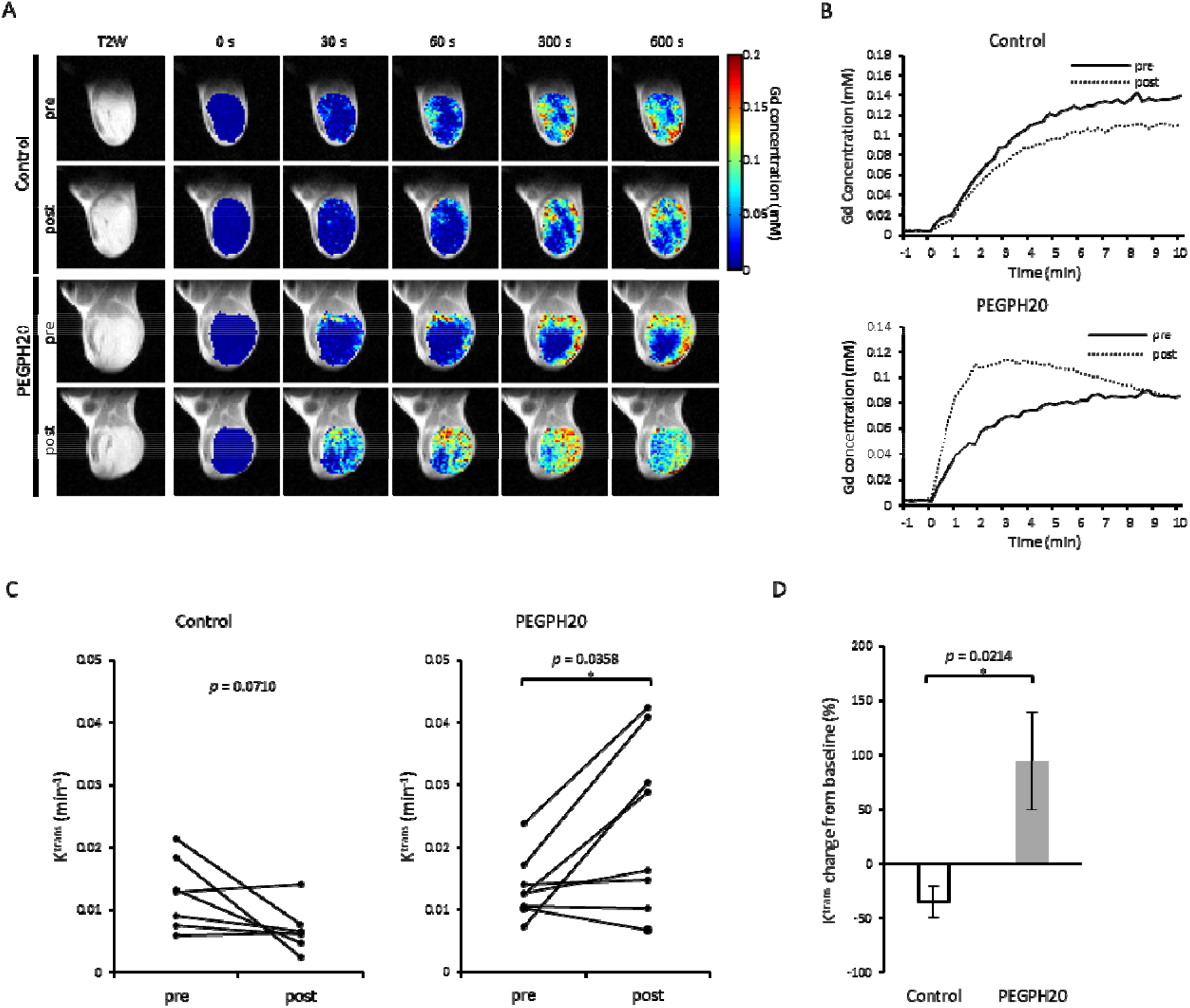
DCE-MRI showed improved intratumor perfusion by PEGPH treatment. DCE-MRI was performed in BxPC3-HAS3 tumor bearing mice treated with control buffer or PEGPH20 (n = 7, n = 8 each group) before (before the first injection on day 0) and after treatment (3 h after the 2nd Ab injection on day3). **A,** Representative images of Gd-DTPA intensity with T2-weighted anatomical image **B,** Representative time-intensity kinetics of Gd-DTPA. **C,** The mean K^trans^ value of each tumor before and after treatment are shown by treatment group. Individual values are shown. **D,** K^trans^ change (%) from the baseline (pretreatment) of each group. Data are shown as mean ± SE. Statistical significance between groups was determined by paired *t*-test for **C** and by Student’s *t*-test for **D.** \**p* < 0.05, \*\**p* < 0.01, \*\*\**p* <0.001 \*\*\*\**p*< 0.0001.

### Effect of PEGPH20 on blood volume

To evaluate the change in patency of vasculature in response to PEGPH20 therapy, blood volume (BV) imaging was also conducted using USPIO and T2 weighted MRI. The local BV (%) in tumors were quantitatively measured from each image of the BxPC3-HAS3 tumors. Figure 4A shows representative anatomical images and BV images at pre-treatment on day 0 and at 30 min post 2nd injection of vehicle or PEGPH20 on day 3, respectively. Distribution of USPIO particles, representing the blood volume, was similarly sparse in both groups before the treatment. However, USPIO particles became globally prominent in images of PEGPH20 treated tumors, while remaining sparse in the vehicle treated tumors. Figure 4B shows the change in BV value of both groups. BV consistently increased in all the 3 mice after PEGPH20 treated group (*p* = 0.0656), while BV decreased in 2 out of 3 mice in the buffer treated group (*p* = 0.3040). The % change in BV was higher in PEGPH20 group relative to the buffer group, although not statistically significant (*p* = 0.0653, Fig.4C). These results suggest the increase in blood volume as a result of the decrease in IFP by PEGPH20 treatment, supporting the findings of improved blood perfusion in the tumor shown in the DCE-MRI studies.

**Figure 4.**
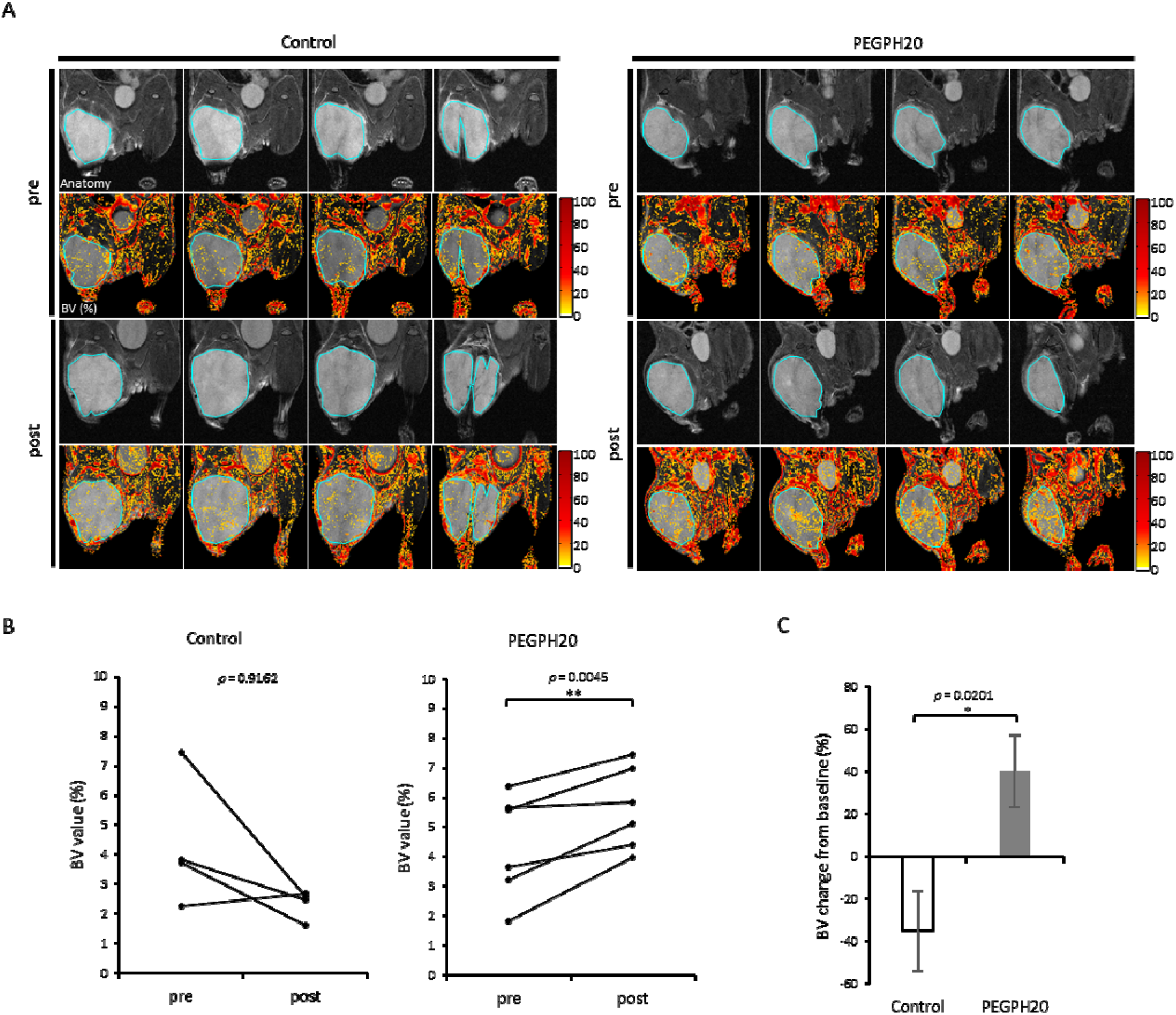
UPIO MRI shows increased blood volume after PEGPH20 treatment. MRI using USPIO was performed in BxPC3-HAS3 tumor bearing mice treated with control buffer or PEGPH20 (n = 3 each group) before (before the first injection on day 0) and after treatment (3 h after the 2nd Ab injection on day3). **A,** Representative imaging of USPIO intensity with T2-weighted anatomical image. Center four slices of each tumor are shown. **B,** Blood volume (%) of region of interest in each tumor (blue line in Fig.3A) before and after treatment are shown by treatment group. Individual values are shown. **C,** Blood volume change (%) from the baseline (pre-treatment) of each group. Data are shown as mean ± SE. Statistical significance between groups was determined by paired *t*-test for **B** and by Student’s *t*-test for **C.**

### Immediate effect of PEGPH20 on intratumor oxygen saturation and total hemoglobin

To better understand the PEGPH20-induced oxygenation in the tumor, photoacoustic (PA) imaging was performed using BXPC3-HAS3 tumors treated with PEGPH20. PA imaging provides a dynamic assessment of changes in oxyhemoglobin (%) and total hemoglobin with an anatomic overlay by calculating from images acquired using near infrared light at two different wavelengths (750 nm and 850 nm). With PA imaging, real-time monitoring of tumor oxygenation is feasible. Figure 5A shows an anatomical image (B mode) followed by PA image before and after the bolus injection of control buffer or PEGPH20 (20 min). The results in the tumor area (purple line, Fig. 5A) were quantified as a change in oxygen saturation of hemoglobin (ΔsO_2_) and % change in total hemoglobin (Hb) and plotted in Figure 5B. Although sO_2_ and total Hb did not show dramatic changes in the control tumor, these parameters rapidly increased during 20 min after the bolus injection of 1mg/kg PEGPH20, which are consistent with improved pO_2_ and increased blood volume. The increase in these parameters after treatment was greater at a higher dose of 10 mg/kg of PEGPH20 (Fig. 5C).

**Fig. 5.**
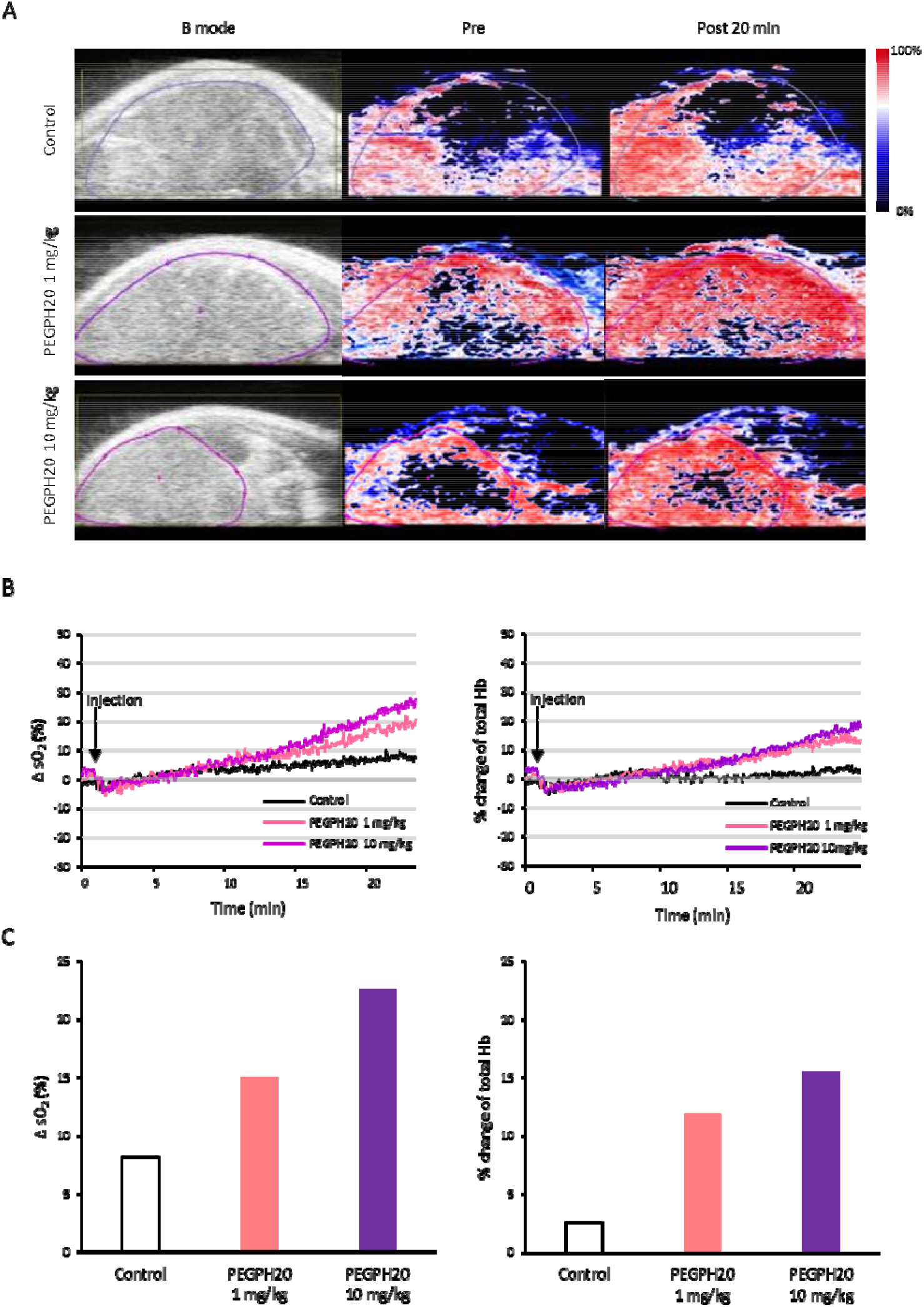
Photoacoustic imaging showed increased oxygen saturation after PEGPH20 injection. Photoacoustic imaging was performed in BxPC3-HAS3 tumor bearing mice treated with control buffer or PEGPH20 for 25 min after the bolus-dose injection (control; n = 1, PEGPH20 1 mg/kg: n=1, PEGPH20 10 mg/kg: n=1). **A,** Anatomical image (B mode) and parametric image of oxygen saturation of hemoglobin (sO_2_) at 20 min after the injection of control buffer (top row), PEGPH20 1 mg/kg (middle row), PEGPH20 10 mg/kg (bottom row). **B,** sO_2_ and total hemoglobin (Hb) of region of interest from the images in Fig. 4A (purple line) are quantified and plotted. **C,** Bar plot of change in sO_2_ (%) and % change of total Hb 20 min after injection.

### Changes in tumor glycolytic metabolism induced by PEGPH20

The change in blood perfusion and oxygenation can influence the metabolic profile of the tumor microenvironment. In particular, increased oxygenation may lead to a metabolic switch from aerobic glycolysis to the TCA cycle as hypoxia is reduced. To examine the resultant metabolic changes after improved perfusion induced by PEGPH20 treatment, we evaluated the pyruvate to lactate flux in vivo by ^13^C MRI using hyperpolarized [1-^13^C] pyruvate. Non-localized 1D spectrum from the entire tumor were acquired continuously for 240 s after the injection of hyperpolarized [1-^13^C] pyruvate and the conversion of pyruvate to lactate was quantitatively evaluated. Representative T2 weighted anatomical scans and kinetics of [1-^13^C] pyruvate and [1-^13^C] lactate of each group are shown in Figure 6A. In Figure 6B, Lac/Pyr values from tumor before and after the treatment was plotted for each mouse. Lac/Pyr ratio increased during 3 days in all the 4 tumors in control group (*p* = 0.0567). The mean Lac/Pyr ratio changed from 0.78 ± 0.09 to 1.10 ± 0.11 after the buffer treatment. In contrast, Lac/Pyr ratio of PEGPH20 treated tumors significantly decreased from 1.18 ± 0.13 to 0.85 ± 0.09 (*p* = 0.0046). The mean changes of Lac/Pyr from baseline value in the buffer and PEGPH20 groups were 44.5 ± 16.4 % and −27.2 ± 3.1 %, respectively, and the difference between groups was statistically significant (*p* = 0.0023, Fig. 6C). The result indicated that improved perfusion leads to improved oxygen availability, resulting in the decreased flux of pyruvate to lactate, presumably from a switch from aerobic glycolysis to the TCA cycle with increased oxygen availability.

**Figure 6.**
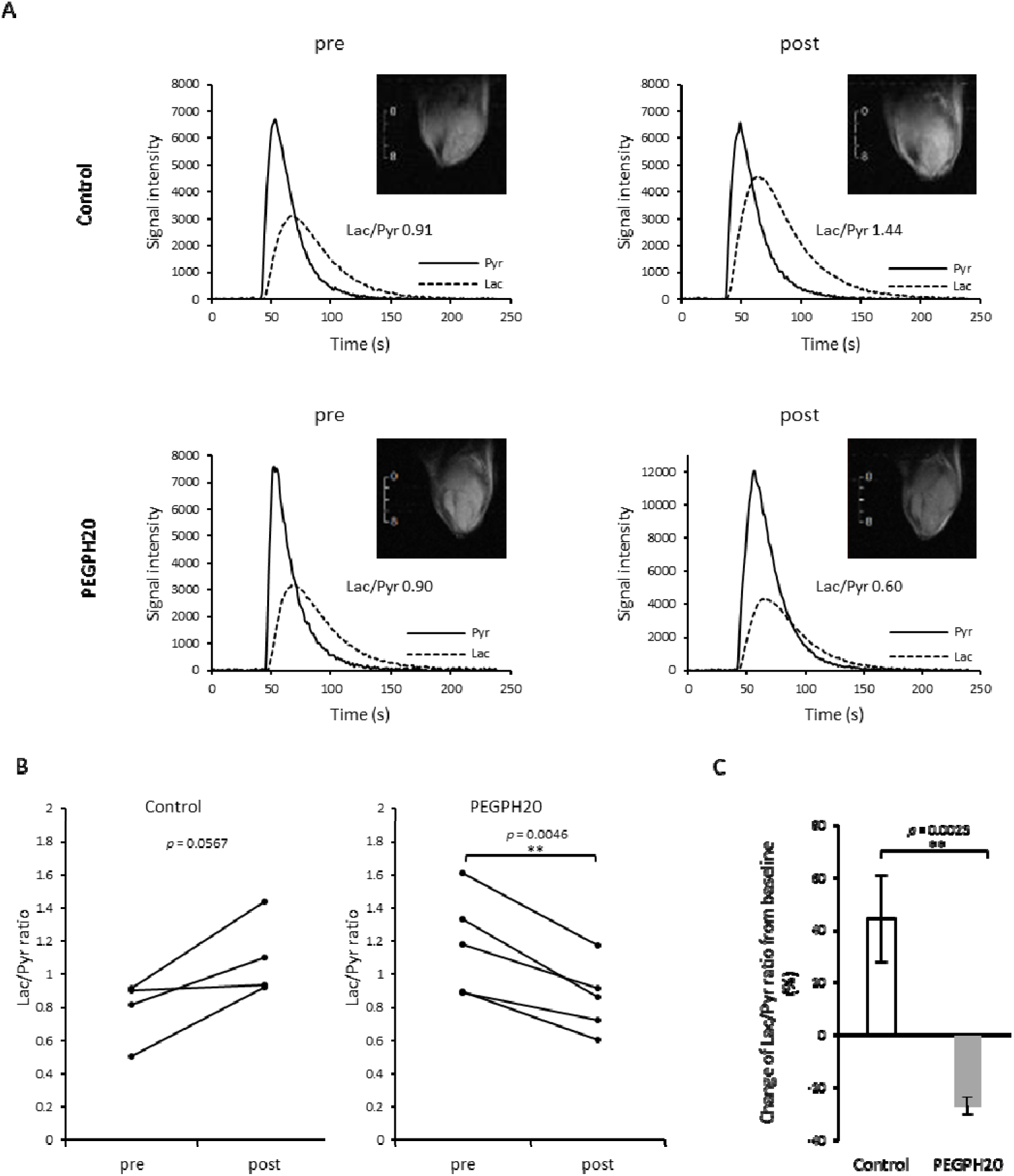
HP [1-^13^C] Pyruvate MRI shows decreased Lac/Pyr ratio after PEGPH20 treatment. HP [1-^13^C] Pyruvate MRI was performed in BxPC3-HAS3 tumor bearing mice treated with control buffer or PEGPH20 (n = 4, n = 5 each group) before (before the first injection on day 0) and after treatment (3 h after the 2nd Ab injection on day3). **A,** Representative kinetics of [1-^13^C] pyruvate and [1-^13^C] lactate and its anatomical ^1^H image. **B,** Lac/Pyr ratio of each tumor before and after treatment are shown by treatment group. Individual values are shown. **C,** Lac/Pyr ratio change from the baseline (pre-treatment) of each group. Data are shown as mean ± SE. Statistical significance between groups was determined by paired *t*-test for **B** and by Student’s *t*-test for **C.** \**p* < 0.05, \*\**p* < 0.01, \*\*\**p* < 0.001 \*\*\*\**p* < 0.0001.

## Discussion

PEGPH20 was originally conceived for use as a combination therapy. By depleting hyaluronic acid, PEGPH20 normalizes the IFP, opening collapsed arterioles and capillaries, increasing blood flow into poorly perfused areas, and improving the outflow within open capillaries (21). By itself this has little effect - PEGPH20 monotherapy did not show a statistically significant clinical benefit in the phase 1 study. The same studies, however, showed PEGPH20 increased the penetration of other drugs (paclitaxel, doxorubicin)(23,37,38) raising the possibility of PEGPH20 as an adjuvant for chemotherapy. However, the phase 3 HALO 301 study, which compared nab-paclitaxel/gemcitabine with and without PEGPH20 in previously untreated patients with stage IV pancreatic cancer, failed to show a clinical benefit of PEGPH20, although a favorable outcome in progression free survival was confirmed in cancer patients treated in an earlier phase 2 trial (27).

Chemotherapy is not the only option for PDAC. Combination use of PEGPH20 with immunotherapy or radiation therapy has not been clinically evaluated. Improved perfusion and local glucose availability in tumor microenvironment has been suggested to be beneficial for immune checkpoint inhibitor therapy in several reports(39,40). One study showed PEGPH20 increased the uptake of anti-programmed death ligand 1 (PD-L1) antibody in HA-accumulating animal models of breast cancer, suggesting that PEGPH20 can sensitize HA-accumulating tumors to anti-PD-L1 immunotherapy (41).

Improved perfusion also contributes to local oxygenation and further radiation sensitization. (42) Previous studies showed that accumulation of HA in tumors was correlated with hypoxia and PEGPH20 treatment decreased the hypoxic fraction when evaluated by histological assessment with pimonidazole staining (23). This indicates that tumor oxygenation improves after PEGPH20 treatment and such a change might be used to predict successful radiation treatment. However, histological assessment for oxygenation is not quantitative and is not a fully reliable method for the detection of hypoxia because it depends both on pimonidazole perfusion and local redox status.(43) In the current study, we employed EPR imaging for the non-invasive and quantitative assessment of pO_2_ following PEGPH20 treatment. The results showed that pO_2_ significantly increased by 4.8 mmHg after treatment with PEGPH20 in BxPC3-HAS3 tumors, while pO_2_ significantly decreased by 2.1 mmHg in tumors treated by the vehicle (Figure 2D). A similar effect was observed within an hour after PEGPH20 treatment by photoacoustic imaging which can record the real-time change in sO_2_, although the scanned area is limited to the superficial tissue which NIR light can reach. Interestingly, the change in sO_2_ was monitored right after a bolus injection of PEGPH20 in our study suggesting an immediate depletion of HA and improvement of perfusion and resultant oxygenation. This result was consistent with the previous report by Thompson et.al showing that PEGPH20 reduced the absolute IFP by 84% in as early as 2 h of a single injection, although the dose was much higher (15 mg/kg) compared to that of our study (1 and 10 mg/kg) (22).

PEGPH20 significantly improved survival times when used in combination with radiation therapy in our xenograft model but only in mice that over-expressed HAS3 (Figure 1). There was no effect on WT mice, which suggests some method of monitoring treatment response will likely be needed if PEGPH20 radiation combination therapy is to proceed to clinical trials. A previous MRI study on PEGPH20 used multiparametric imaging including glycosaminoglycan chemical exchange saturation transfer (GagCEST) and apparent diffusion coefficient mapping to quantify changes in the tumor microenvironment in mice using a 14 tesla scanner (44). A decreased GagCEST effect and increased ADC value suggested improved tumor perfusion after PEGPH20 treatment. The need for a high field to quantify the GagCEST effect may prove problematic for future clinical studies. In the current study, we focused on finding useful imaging biomarkers that can be acquired at the lower field scanners (3 tesla) available for human subject research. The increase in K^trans^ after PEGPH20 treatment is indirect evidence of decreased IFP. Similarly, the increase in blood volume measured by photoacoustic imaging is direct evidence of improved patency of vasculature. The results are consistent with a previous preclinical study showing that increased K^trans^ was detected 24hr after PEGPH20 injection in an animal model (38) and a clinical study also showing increased K^trans^ after treatment in majority of tumors including pancreatic cancer in 11 patients (37). These imaging biomarkers may be useful to predict the effect of PEGPH20 enhancing the efficacy of other chemotherapeutics because a higher level of HA depletion serum marker in cancer patients treated with nab-paclitaxel/gemcitabine was associated with better survival in the previous phase 2 clinical trial (45).

The effectiveness of PEGPH20 as a combination for immunotherapy depends not only on increasing immune infiltration but also the effect PEGPH20 has on the entire tumor microenvironment, in particular whether the tumor can sustain the metabolic activity necessary for immune activity. The effect of PEGPH20 on tumor metabolism is controversial. There are a few studies reporting an association between tumor cell metabolism and HA in tumor microenvironment. In breast cancer, HA production shifts metabolic programs toward acceleration of glycolysis, (46) which is regulated by HIF-1 under the control of the hexosamine biosynthetic pathway flux.(47) Consistently, it has been reported that antagonizing hyaluronan-CD44 interactions with HA oligomers inhibits the assembly of MCT1 and MCT4 complexes in the plasma membrane, and consequently suppresses lactate efflux from breast carcinoma cells (48). In addition, one Phase 1 trial of PEGPH20 evaluating ^18^F-FDG-PET/CT showed that 6 out of 9 evaluable patients exhibited decrease in standardized uptake value (SUV), suggesting an upregulation of glycolysis (37). Similarly, one report showed that treatment of cancer cells with hyaluronidase triggers an increase in glycolysis through degradation of TXNIP, which promotes internalization of the glucose transporter GLUT1 (49). In the current study, ^13^C pyruvate MRI showed PEGPH20 treatment decreased pyruvate to lactate flux in BxPC3-HAS tumors. We observed a similar metabolic shift in a mouse xenograft model of SCC7 tumors treated with sunitinib targeting for vascular normalization (31). This suggests that glycolytic flux decreases when perfusion and oxygenation improves in tumor microenvironment, likely due to decreased activity in the HIF-1 pathway that upregulates glucose transporter-1 and pyruvate dehydrogenase kinase (which inhibits the activity of pyruvate dehydrogenase) as well other enzymes in the glycolytic pathway (50).

In conclusion, by using non-invasive multimodal imaging modalities, we could confirm a radio-sensitizing effect of PEGPH20 in a xenograft model. The effect is attributed to a change in the tumor microenvironment where oxygenation is improved and pyruvate to lactate flux is decreased by PEGPH20. These factors together likely account for the enhanced radiation efficacy after PEGPH20 treatment. The imaging modalities used in this study comprehensively clarify the hemodynamic and metabolic changes induced by treatments modifying extracellular matrix in tumor microenvironment. Further, these pre-clinical imaging data suggest that ECM degradation with PEGPH20 already tested in humans can be a strategy to achieve radiasensitization for improved treatment efficacy and outcome.

## Author Contributions

Tomohiro Seki: Investigation, Methodology, Yu Saida: Investigation, Methodology, Writing – original draft, Shun Kishimoto: Investigation, Conceptualization, Methodology, Supervision, Writing-original draft, Jisook Lee: Resources, Methodology, Yasunori Otowa: Investigation, Methodology, Kazutoshi Yamamoto: Investigation, Gadisetti VR Chandramouli: Formal Analysis, Investigation, Software, Nallathamby Devasahayam: Resources, Investigation, James B. Mitchell: Project Administration, Supervision, Funding acquisition, Jeffrey R. Brender: Conceptualization, Formal Analysis, Software, Writing – review & editing, Murali C. Krishna: Conceptualization, Project Administration, Methodology, Supervision, Funding acquisition

